# Rapid Differentiation of Epithelial Cell Types in Forensic Samples Using Intrinsic Fluorescence and Morphological Signatures

**DOI:** 10.1101/245159

**Authors:** Emily R. Brocato, M. Katherine Philpott, Catherine C. Connon, Christopher J. Ehrhardt

## Abstract

Establishing the tissue source of epithelial cells within a biological sample is an important capability for forensic laboratories. In this study we used Imaging Flow Cytometry (IFC) to analyze individual cells recovered from buccal, contact epithelial, and vaginal samples that had been dried between 24 hours and more than eight weeks. Measurements capturing the size, shape, and fluorescent properties of cells were collected in an automated manner and then used to build a multivariate statistical framework for differentiating cells based on tissue type. Results showed that cell populations from the three tissue types could be differentiated using a Discriminant Function plot of IFC measurements. Epidermal cells were distinguished from vaginal and buccal cells with an average classification accuracy of ~94%. Ultimately, cellular measurements such as these, which can be obtained non-destructively, may provide probative information for many types of biological samples and complement results from standard genetic profiling techniques.

## Introduction

Characterizing the type of cells present in biological evidence and, therefore, the tissue they originated from within the body, can assist with crime reconstructions and downstream DNA profiling methods. Traditionally, caseworking methods for determining tissue source are based on microchemical and/or enzymatic reactions targeted toward proteins within bodily fluids, which have limited sensitivity and/or specificity. Recently, there has been considerable research into biomolecular markers for tissue identification. These include mRNA transcripts (1), micro-RNAs (2,3), proteomics (4), and DNA methylation patterns (5). Although promising, the specificity of many of these systems is still being investigated and interpretation can require complex bioinformatic workflows.

In contrast, few forensic techniques have utilized morphological or intrinsic biochemical differences to differentiate between cells from different tissues, particularly epithelial cells. This is likely due to the laborious nature of microscopic characterizations or the need for tissue-specific antibody probes which have limited success on dried or compromised samples (6,7). One potential strategy to address these challenges is the use of Imaging Flow Cytometry (IFC). In IFC, conventional flow cytometry analysis whereby the optical properties of individual cells are interrogated with lasers at set wavelengths, is combined with fluorescence and bright field imaging of those same cell events. IFC is routinely used in biomedical and clinical research for identification of unusual cell types as well as high resolution surveys of sub-cellular processes (8). The primary advantage of IFC over conventional microscopic analysis is that images of single cells are collected in a high throughput manner (as many as hundreds per second) and at multiple fluorescence channels simultaneously. The resulting multivariate data streams can therefore be used to compare profiles between individual cells or between larger populations. For forensic applications, another potential advantage of IFC is that it is an inherently nondestructive technique, with the possibility of collecting all cells after analysis for DNA profiling or other biological characterizations.

In this study we tested whether IFC could be used to differentiate epithelial cells from three separate tissue sources: buccal, touch epidermal, and vaginal. Identifying the presence of one or more of these cell types in a biological sample when combined with DNA profiling results may be useful when investigating claims of sexual assault, particularly without ejaculation. Additionally, because DNA yield has been observed to systematically vary between epidermal cells and other types of epithelial tissue (9), determining the presence and relative quantities of each cell type can help direct downstream DNA profiling efforts. We conducted an initial IFC survey of each epithelial cell type by analyzing existing biological specimens from a forensic sample repository consisting of ten donors per cell type. To assess the robustness and consistency of IFC signatures against samples approximating those that would be encountered during forensic casework, each of the three cell types were collected from multiple donors and were “aged” for different amounts of time prior to analysis, ranging from ~24 hours to more than eight weeks.

## Methods

### Sample Collection and Preparation

Buccal samples were obtained pursuant to VCU-IRB approved protocol ID#HM20000454_CR3. Ten volunteers were asked to swab the inside of cheek for 30 seconds. For dried samples, swabs were left to dry for 1 hour to 6 days. Dried and fresh swabs were processed in the same manner. Contact samples were obtained pursuant to VCU-IRB approved protocol ID# HM20000454_CR3. Ten volunteers (six of whom were buccal cell donors) were asked to hold/rub a conical tube (P/N 229421; Celltreat Scientific) for five minutes to deposit cells. For dried samples, tubes were left out for 24 hours to 5 days to dry before collecting cells. Cells were collected from the surface with 1 sterile, pre-wetted swab, and 1 sterile, dry swab. Ten vaginal samples were obtained pursuant to VCU-IRB approved protocol ID#HM20002931_Ame2. Volunteers were asked to swab the inside of the vaginal cavity, and swabs were dried and stored at room temperature until analysis. Storage times ranged from 48 hours to approximately eight weeks. All collection swabs were eluted in 1 mL of 1x Cell Staining Buffer (P/N 420201; Biolegend), and gently vortexed for 10 seconds. Samples were centrifuged at 1500 × g at 4°C for 5 minutes. The supernatant was discarded, and the cell pellets were dissolved in 100 uL of 1x Cell Staining Buffer for imaging flow cytometry.

A list of all donor samples used in this study and their respective drying times are provided in Table S1.

### Imaging Flow Cytometry and Statistical Analysis

All samples were analyzed using an Amnis^®^ Imagestream X Mark II (EMD Millipore, Burlington MA) equipped with 405nm, 488nm, 561nm, and 642nm lasers. Laser voltages for all tests were set at 120mW, 100mW, 100mW and 150mW, respectively. Images of individual events were captured in five detector channels labeled: 1 (430-505nm), 2 (505-560nm), 3 (560-595nm), 5 (640-745nm), and 6 (745-780nm). Channel 4 was used to capture Brightfield images. Magnification was set at 40x and autofocus was enabled so that the focus varied with cell size. Examples of cell images collected across multiple wavelengths are provided in Figure S1. Raw image files (.rif) were then imported into IDEAS^®^ Software (EMD Millipore). Display Width and Display Height were changed to 120×120 pixels for each image. The ‘Shape Change Wizard’ option in Ideas was used to gate focused cells on a Gradient RMS_M04Ch04 × Normalized Frequency histogram. Once focused cells were selected, single cells were gated on an Area_M04 × Aspect Ratio_M04 scatterplot. Data for individual cell events were collected for 17 different features: area, aspect ratio, aspect ratio intensity, contrast, intensity, mean pixel, median pixel, max pixel, raw max pixel, raw min pixel, length, width, height, brightness detail intensity (‘R3’ pixel increment), raw centroid X, raw centroid Y, and circularity. These feature measurements were collected across multiple detector channels (i.e., fluorescence and brightfield wavelengths) with the exception of measurements that could only be determined from brightfield images such as centroid X/Y and circularity. This yielded a total of 88 measurements/variables collected for each cell. Feature data for individual cells was then exported into a Microsoft Excel worksheet. Cell yield varied across each of the study samples but did not appear to be correlated with tissue type, drying time, or individual donor. Most cell populations yielded between 200 and 400 cell images with nine samples providing between 80 and 200 images.

IFC measurement values were then imported into SPSS v23 (IBM, Inc. Chicago, IL). Differences in mean values between the three cell types were tested using a one-way ANOVA analysis with a Tukey HSD post-hoc test performed in SPSS. Next, multivariate differences among the three cell type groups were analyzed using a Discriminant Function Analysis (DFA) based on the within-group covariance matrix. We initially compared results from direct analysis of IFC measurements and those obtained from transforming the data first into principal components (PCs) and then conducting DFA on the PC scores. We found that the latter approach led to less differentiation in the canonical variate plot and poorer classification accuracy and thus used direct analysis of raw measurements. Different combinations of variables were initially tested based on their impact on group separation in the canonical variate plot and classification accuracy. Inclusion of all variables in the analysis resulted in the greatest degree of separation in the canonical variate plot and the highest rate of accurate classifications.

## Results and Discussion

We first tested whether IFC could be used to distinguish cells from the three different epithelial tissue sources. During sample processing and image collection, we noted some general qualitative differences between images from each of the three cell types. For example, nuclei were routinely observed in brightfield images of buccal cells and vaginal cells (e.g., Images 1507, 1796, respectively, Figure 1), while they were rarely observed in contact epithelial cell images. Buccal and vaginal cells were generally larger in size, >40 μm compared to contact cells, which were ~20-50 μm although we noted some size overlap between cell sources. This could be due in part from the folding or degradation of buccal and vaginal cells during drying or sampling prior to IFC. Contact cells generally exhibited higher contrast features in brightfield images compared to buccal or vaginal cells.

**Figure 1.**
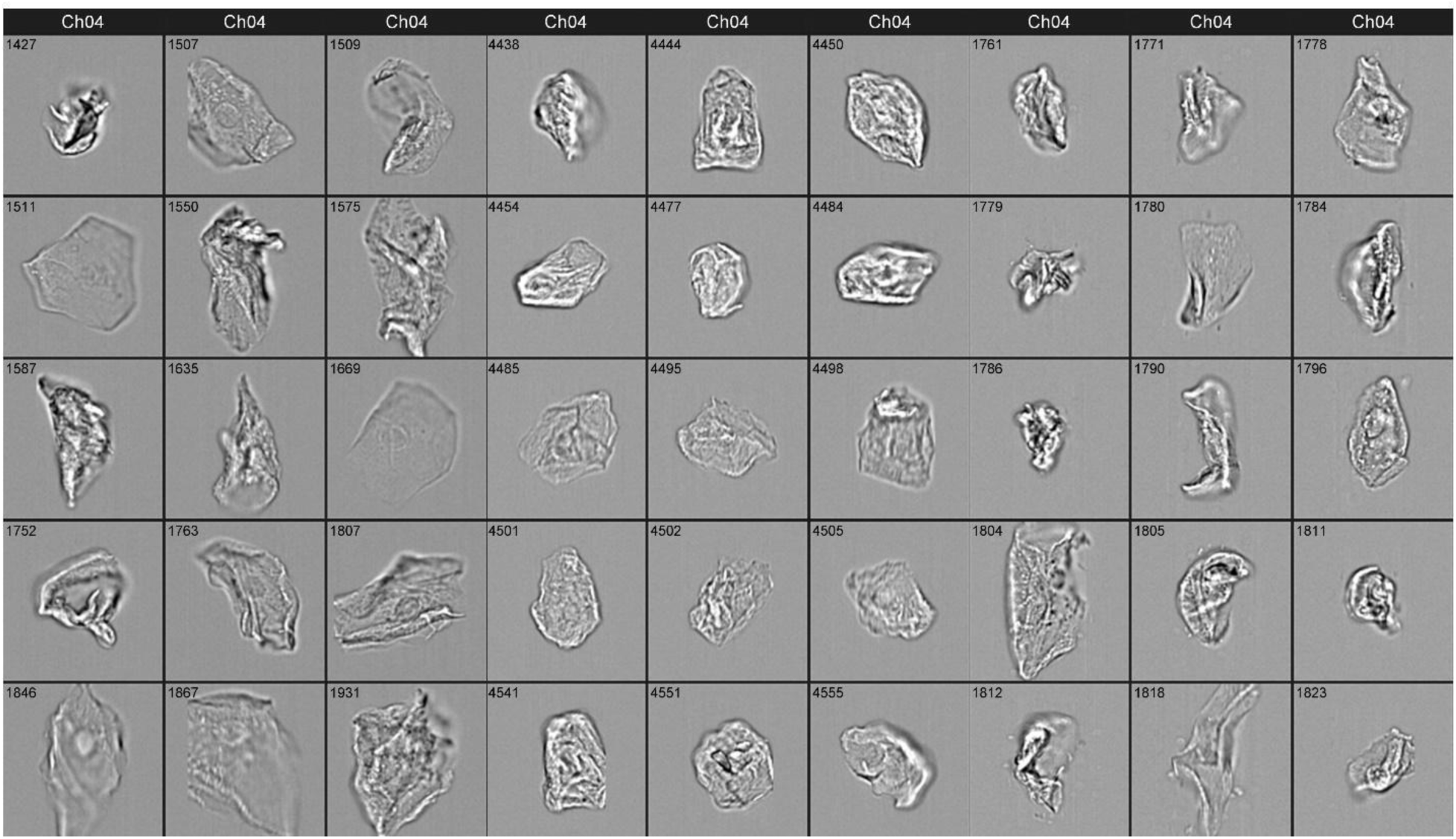
Top-IFC brightfield image gallery for buccal cells (columns 1-3), contact epithelial cells (columns 4-6), and vaginal cells (columns 7-9). Each image frame is 50 μm × 50 μm. Object identifier are included with each image.

For the 264 pairwise comparisons between group means (88 variables and three sample groups), only 42 yielded p-values greater than 0.01, with the vast majority showing p values less than 0.0001 (Table S2). Of note were differences in means for circularity (7.8 contact, 4.1 buccal, 4.3 vaginal), intensity (e.g., in 430-505nm channel 3×10^5^ RFU contact, 6×10^4^ RFU buccal, 5×10^4^ RFU vaginal), and brightness detail (e.g., in 403-505 nm channel 1×10^4^ RFU contact, 9×10^3^ buccal, 7×10^3^ RFU vaginal). However, the range of values for each cell group showed a high degree of overlap across the three cell types (Boxplots in Figure S2). Similarly, most variables showed large standard deviations for each cell type, with coefficients of variation for individual measurements ranging from ~20% to more than 280%.

In order to determine whether the observed variation in IFC measurements could be used to differentiate cell types, we employed Discriminant Function Analysis (DFA) as a supervised multivariate technique to model variation between groups. In DFA, linear combinations of the original variables are constructed (i.e., canonical variates) such that the variation between user-defined sample groups is maximized and within group variation is minimized. DFA is a well-established technique with demonstrated applications for other forensic signature systems (10–12) For this dataset, the primary advantages of DFA are that differences in measurement scales across variables do not impact the analysis and it is relatively robust to non-normally distributed data (13). Additionally, the canonical variates generated with DFA can be used to classify individual samples into one of the user-defined groups. For this study we used DFA to initially examine multivariate differences between groups. A DFA plot of all IFC measurements from all three cell types showed distinct separation between buccal, contact epithelial, and vaginal cell populations (Figure 2). Multivariate differences between groups were statistically significant, Wilk’s Lambda= 0.114, p<0.001. Some overlap is observed among the sample groups on the DFA plot, in particular between buccal and vaginal cell groups. A leave-one-out (LOO) classification on individual cell images for each of the three groups and all 30 donor cell populations showed an classification accuracy of ~90%.

**Figure 2.**
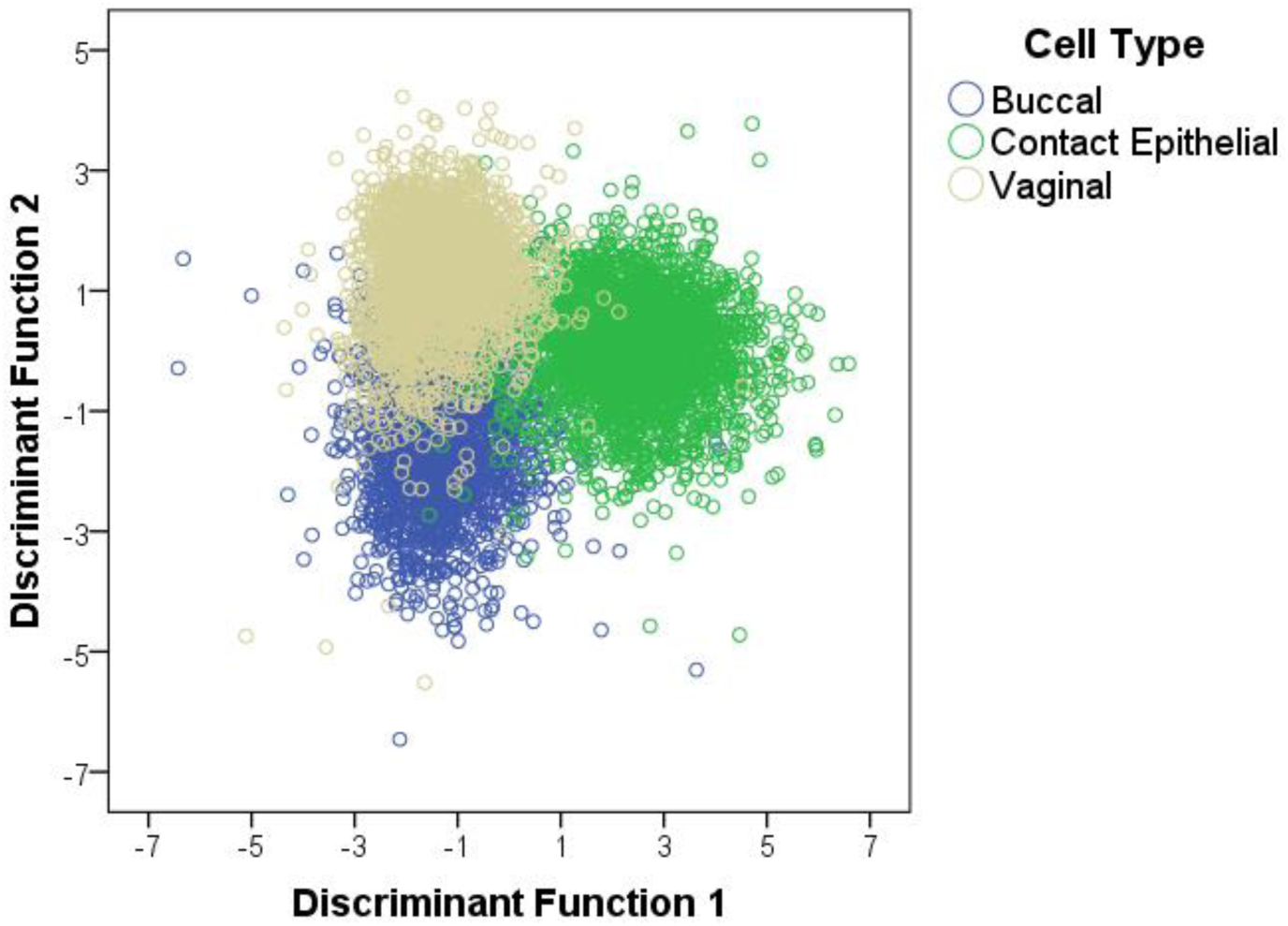
Discriminant Function Analysis of epithelial cells from three tissue sources using IFC variables. The first discriminant function (x-axis) accounted for ~74% of the between group variation and the second discriminant function (y-axis) accounted for ~26%.

Next we used DFA-based algorithms to classify entire donor cell populations into one of the three cell groups in a blinded fashion to determine the accuracy and robustness of this approach for identifying cell types from an unknown forensic sample. This was accomplished by withholding a given donor cell population from the DFA and classifying each cell image into one of the three epithelial cell types based on information from the remaining contributor cell populations. Classification results for cell populations against three tissue types are given in Table 1. In general, epidermal cells showed the highest classification accuracy (89%) with six of the ten donor cell populations having accuracies over 90%. Only one cell population, P22, was below 80%. Buccal and vaginal cell populations yielded lower classification rates, 77% and 75% respectively (Table 1). Interestingly, classification rates were highly variable across individual cell populations for these two groups, with buccal cells ranging between 55% and 90% and vaginal cells ranging between 29% and 97%.

**Table 1.** Classification Test of Buccal, Epidermal and Vaginal Cells

| Cell Type | Contributor ID | Predicted Cell Type |  |  | Classification % |
| --- | --- | --- | --- | --- | --- |
|  |  | Buccal | Epidermal | Vaginal |  |
| Buccal | L49 | 261 | 15 | 32 | 85 |
| Buccal | I66 | 244 | 2 | 70 | 77 |
| Buccal | C58 | 232 | 12 | 17 | 89 |
| Buccal | R47 | 149 | 16 | 0 | 90 |
| Buccal | 5001 | 261 | 3 | 88 | 74 |
| Buccal | B21 | 330 | 22 | 13 | 90 |
| Buccal | N08 | 273 | 28 | 3 | 90 |
| Buccal | Y60 | 252 | 26 | 82 | 70 |
| Buccal | Z32 | 244 | 31 | 90 | 67 |
| Buccal | 5034 | 226 | 27 | 155 | 55 |
| **Total** | **--** | **2472** | **182** | **550** | **77** |
| Epidermal | Q17 | 4 | 191 | 8 | 94 |
| Epidermal | Z32 | 20 | 178 | 5 | 88 |
| Epidermal | Y60 | 5 | 192 | 5 | 95 |
| Epidermal | K36 | 0 | 421 | 12 | 97 |
| Epidermal | P22 | 34 | 222 | 107 | 61 |
| Epidermal | S95 | 3 | 79 | 2 | 84 |
| Epidermal | I66 | 14 | 309 | 13 | 92 |
| Epidermal | L49 | 2 | 348 | 1 | 99 |
| Epidermal | R47 | 51 | 251 | 0 | 83 |
| Epidermal | N08 | 5 | 277 | 21 | 91 |
| **Total** | **--** | **138** | **2468** | **174** | **89** |
| Vaginal | 2368 | 200 | 18 | 160 | 42 |
| Vaginal | 4017 | 11 | 13 | 288 | 92 |
| Vaginal | 1022 | 46 | 13 | 298 | 84 |
| Vaginal | 1028 | 93 | 2 | 37 | 29 |
| Vaginal | 1031 | 1 | 9 | 357 | 97 |
| Vaginal | 5020 | 40 | 0 | 128 | 76 |
| Vaginal | 5021 | 1 | 26 | 110 | 80 |
| Vaginal | 5005 | 24 | 36 | 108 | 64 |
| Vaginal | 4502 | 14 | 0 | 186 | 93 |
| Vaginal | 4504 | 75 | 0 | 227 | 75 |
| **Total** | **--** | **505** | **117** | **1899** | **75** |

In an attempt to improve the classification accuracy for each cell type, individual cell populations were also tested with two-group classification schemes where one tissue group was excluded completely from the analysis, i.e., buccal cells against epidermal cells; vaginal cells against epidermal cells; and buccal cells against vaginal cells (Tables 2–4 respectively). Simplified classification schemes could be run subsequent to the original classification to help identify samples assigned to one of the closely related sample groups, i.e., a cell image classified as a buccal cell in the three group DFA could then be run against a two group DFA containing only buccal and vaginal cells. Tiered or successive DFA analyses have been described for other types of forensic samples (10,14). Additionally, two group comparisons could approximate caseworking scenarios in which one of the epithelial cell types could be ruled out *a fortiori* for an unknown cell population. Results from two-group DFA showed that buccal and epidermal cell populations could be differentiated with the highest accuracy (~94%). The lowest classification rate of individual donor cell populations in this comparison was 83% (P22, Epidermal) with the majority of cell populations exhibiting classification rates of 95% or higher (Table 1). The vaginal-epidermal cell classifications showed comparable results with an overall classification rate of ~91%. Two individual cell populations in this scheme exhibited markedly lower success rates, e.g., P22 epidermal 67% and 5005 vaginal 55%. However, the remaining cell populations had classification rates >80% with the majority >95% (Table 2). The poorest differentiation was observed between buccal and vaginal cells with an overall classification accuracy of 79% (Table 3) and multiple donor cell populations below 60% accuracy (e.g., 5034, Buccal; Z32 Buccal; 2368 Vaginal).

**Table 2.** Classification Test of Buccal vs. Touch Epidermal Cells

| Cell Type | Contributor ID | Predicted Cell Type |  | Classification % |
| --- | --- | --- | --- | --- |
|  |  | Buccal | Contact Epithelial |  |
| Buccal | L49 | 287 | 20 | 93 |
| Buccal | I66 | 308 | 7 | 98 |
| Buccal | C58 | 232 | 28 | 89 |
| Buccal | R47 | 140 | 24 | 85 |
| Buccal | 5001 | 348 | 3 | 99 |
| Buccal | B21 | 197 | 4 | 98 |
| Buccal | N08 | 135 | 5 | 96 |
| Buccal | Y60 | 175 | 21 | 89 |
| Buccal | Z32 | 173 | 28 | 86 |
| Buccal | 5034 | 234 | 10 | 96 |
| **Total** | **--** | **2229** | **150** | **94** |
| Epidermal | N08 | 6 | 297 | 98 |
| Epidermal | R47 | 31 | 271 | 90 |
| Epidermal | L49 | 2 | 349 | 99 |
| Epidermal | I66 | 8 | 328 | 98 |
| Epidermal | S95 | 3 | 81 | 96 |
| Epidermal | P22 | 61 | 302 | 83 |
| Epidermal | K36 | 0 | 433 | 100 |
| Epidermal | Y60 | 2 | 200 | 99 |
| Epidermal | Z32 | 19 | 184 | 91 |
| Epidermal | Q17 | 2 | 201 | 99 |
| **Total** | **--** | **134** | **2646** | **95** |

**Table 3.** Classification Test Vaginal vs. Touch Epidermal Cells

| Cell Type | Contributor ID | Predicted Cell Type |  | Classification % |
| --- | --- | --- | --- | --- |
|  |  | Vaginal | Contact Epithelial |  |
| Vaginal | 2368 | 312 | 66 | 82 |
| Vaginal | 4017 | 283 | 29 | 91 |
| Vaginal | 1022 | 301 | 56 | 84 |
| Vaginal | 1028 | 113 | 19 | 86 |
| Vaginal | 1031 | 363 | 4 | 99 |
| Vaginal | 5020 | 166 | 2 | 99 |
| Vaginal | 5021 | 123 | 14 | 90 |
| Vaginal | 5005 | 93 | 75 | 55 |
| Vaginal | 4502 | 200 | 0 | 100 |
| Vaginal | 4504 | 302 | 0 | 100 |
| **Total** | **--** | **2256** | **265** | **89** |
| Epidermal | Q17 | 8 | 195 | 96 |
| Epidermal | Z32 | 10 | 193 | 95 |
| Epidermal | Y60 | 5 | 196 | 98 |
| Epidermal | K36 | 23 | 410 | 95 |
| Epidermal | P22 | 119 | 244 | 67 |
| Epidermal | S95 | 4 | 80 | 95 |
| Epidermal | I66 | 9 | 327 | 97 |
| Epidermal | L49 | 2 | 349 | 99 |
| Epidermal | R47 | 0 | 302 | 100 |
| Epidermal | N08 | 19 | 284 | 94 |
| **Total** | **--** | **199** | **2580** | **93** |

**Table 4.** Classification Test of Buccal vs. Vaginal Cells

| Cell Type | Contributor ID | Predicted Cell Type |  | Classification % |
| --- | --- | --- | --- | --- |
|  |  | Buccal | Vaginal |  |
| Buccal | L49 | 291 | 16 | 95 |
| Buccal | I66 | 249 | 66 | 79 |
| Buccal | C58 | 256 | 4 | 98 |
| Buccal | R47 | 164 | 0 | 100 |
| Buccal | 5001 | 230 | 121 | 66 |
| Buccal | B21 | 195 | 6 | 97 |
| Buccal | N08 | 136 | 4 | 97 |
| Buccal | Y60 | 148 | 48 | 76 |
| Buccal | Z32 | 109 | 92 | 54 |
| Buccal | 5034 | 52 | 192 | 24 |
| **Total** | **--** | **1830** | **549** | **77** |
| Vaginal | 2368 | 256 | 122 | 32 |
| Vaginal | 4017 | 16 | 296 | 95 |
| Vaginal | 1022 | 26 | 331 | 93 |
| Vaginal | 1028 | 55 | 77 | 58 |
| Vaginal | 1031 | 0 | 368 | 100 |
| Vaginal | 5020 | 39 | 131 | 77 |
| Vaginal | 5021 | 1 | 139 | 99 |
| Vaginal | 5005 | 20 | 152 | 88 |
| Vaginal | 4502 | 11 | 194 | 95 |
| Vaginal | 4504 | 59 | 249 | 81 |
| **Total** | **--** | **483** | **2059** | **81** |

Overall, the relatively high classification rates of epidermal cells against buccal cells and epidermal cells against vaginal cells (>90%) suggests that systematic differences in morphological and/or optical properties measured by IFC can be used to distinguish between epithelial cell types in these two comparison schemes. Further, measurement values can potentially be used to construct an analysis framework for characterizing unknown cell populations into one of these three sample groups. The observed variation between sloughed epidermal cells and buccal/vaginal cells is consistent with the intrinsic biochemical, structural, and morphological differences for cells originating from each tissue source. For example, shed epidermal cells are derived from the stratum corneum and characterized by a high degree of keratinization with few if any organelles and little intracellular DNA owing to the apoptotic processes occurring as cells migrate from the basal to the upper layers of the epidermis (15). In contrast, buccal and vaginal cells are derived from less stratified epithelial tissue and may be only partially keratinized or unkeratinized. Although no studies to date have explicitly surveyed cellular differences between these three tissue sources using fluorescence signatures, previous work has shown that changes in cellular autofluorescence can be used to differentiate layers of epidermal tissue with different intracellular components (e.g., keratin, tryptophan, FAD) (16,17). Additionally, the morphological and size differences detected with IFC (e.g., area and circularity measurements) are consistent with histological context of each cell type, i.e., shed epidermal cells hexagonal and ~20-50 μm, while buccal and vaginal cells are typically >40 μm with elongated shapes (18,19).

The overlap between cell sources shown in Figure 2 and misclassifications of individual cells images, could be due to a number of factors. First, some similarities in fluorescence and/or morphological attributes are expected, particularly for buccal and vaginal cells given that both are derived from non-keratinized epithelial tissue. This is consistent with poorer classification rates of buccal/vaginal cells relative to buccal-epidermal and vaginal-epidermal (Table 3 vs. Tables 1–2 respectively). Second, cell populations in this data set represent a wide range of drying/exposure times prior to sampling and analysis. Levels of intrinsic fluorescence are likely to change with time owing to the degradation of cellular components such that specimens with longer periods of environmental exposure may be harder to distinguish from each other. Although there were no clear relationships between exposure time and position on the DFA plot (Figure 2) or classification accuracy (Tables 1–3), this should be explicitly tested with future studies. Another factor that could be contributing to misclassifications is inter-individual variation. Previous work from our group has shown that autofluorescence signatures in shed epidermal cells can vary between contributors, likely owing to the presence of exogenous materials associated with the cell (20). Cell populations from different contributors of the same tissue type (epidermal or buccal) and drying time (24 or 48 hours, respectively) showed distinct separation in a preliminary DFA (Figure 3). It is also possible that the number of donor cell populations in this study did not adequately capture the full range of morphological and/or fluorescence variation that exists between contributors. Increasing the number of unique donor cell populations in the reference/comparison dataset will likely help to isolate any tissue-specific signatures that are present. Nevertheless, contributor-specific variation in IFC measurements is a potentially promising avenue of future research for this technique, particularly how it might be used for estimating the number of individual cell populations in a biological sample and/or facilitating front-end cell separation in a DNA profiling workflow.

**Figure 3.**
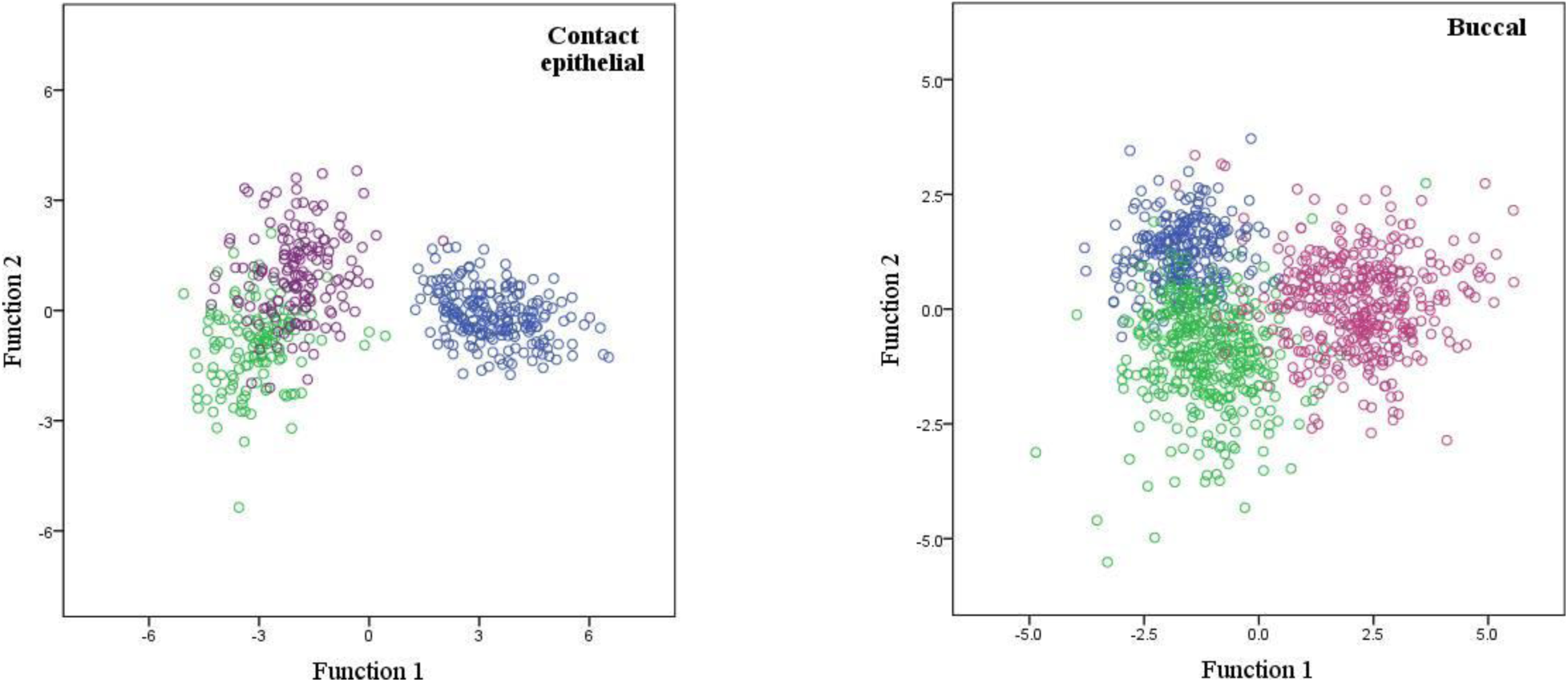
Discriminant function analysis of cell populations derived from three different contributors for contact epithelial (left) and buccal (right) tissue sources. Contact epithelial cell samples were dried for 24 hours at room temperature and buccal cells were dried for 48 hours at room temperature prior to analysis.

## Conclusion

Our goal with this study was to conduct an initial assessment of high-throughput fluorescence and morphological analysis and its potential applications for characterizing epithelial cell types in an unknown biological sample. High-throughput, single cell measurements combined with a multivariate classification framework were used to distinguish epidermal cells from other epithelial cell sources across a range of drying times with an overall high degree of accuracy. Although a range of factors may contribute to morphological or optical properties in any given sample (e.g., individual-specific signatures and degradation time), these results suggest that multivariate approaches may be used to extract tissue-specific signatures from biological samples. Future work should test alternative classification methods such as machine learning algorithms as well as different combinations of cellular measurements to maximize classification accuracy particularly for cell types with similar biochemical and physical properties but different source tissue, and eventually establish cell count thresholds for inferring the presence and/or proportions of one or more cell types in a biological sample. Lastly, we note that the fluorescence and morphological signatures identified here could be detected and analyzed using other microscopy setups and open source software platforms (e.g., (21)) which may facilitate its use in caseworking laboratories.

## Acknowledgements

The authors gratefully acknowledge Julie Farnsworth for providing technical assistance for this project.

